# Evaluating an In Vivo Oxidative Stress sensor in cystic fibrosis rat epithelial tissues

**DOI:** 10.1101/2025.11.25.690596

**Authors:** Nathan Rout-Pitt, Serena Barnes, Nicole Reyne, Alexandra McCarron, Martin Donnelley, Roman Kostecki, Erik Noschka

**Affiliations:** Robinson Research Institute, University of Adelaide, Adelaide, South Australia, Australia; Adelaide Medical School, University of Adelaide, Adelaide, South Australia, Australia; School of Animal and Veterinary Science, University of Adelaide, Adelaide, South Australia; Respiratory and Sleep Medicine, Women’s and Children’s Hospital, Adelaide, South Australia, Australia; Sensrz, Adelaide, South Australia, Australia

**Author notes:** Co-senior author (Roman Kostecki, Erik Noschka). Co-first author (Nathan Rout-Pitt, Serena Barnes).

**Keywords:** Cystic Fibrosis, oxidative stress, epithelial, sensor

## Abstract

We have developed a novel biosensing device that can detect the real-time, dynamic state of *in vivo* oxidative status (IVOS) in living systems. Oxidative stress is a well-established condition in CF animal models and humans. Elevated oxidative stress conditions are associated with excessive inflammatory responses from neutrophils that result in fibrotic tissue formation. As such, numerous clinical and preclinical studies suggest that elevated oxidative stress, combined with the heightened pro-inflammatory milieu observed in CF phenotypes, likely increases susceptibility to recurrent infection–inflammation cycles. We aimed to use the IVOS sensor to characterise oxidative environmental differences in wildtype, CF *Phe508del* and CFTR knockout rat epithelial tissues including the lungs, trachea, nasal mucosa, and oesophagus. Our results revealed no significant differences in the baseline redox balance between wildtype, Phe508del and knockout rat tissues, however when we looked at short term fluctuations in redox status using the ratio of the root mean squared of successive differences (RMSSD) to the normalised mean IVOS arbitrary units, we saw significantly elevated fluctuations in the redox status in the bronchi of Phe508del rats compared to wildtype rats, indicating a lower steady-state oxidative environment combined with large transient oxidative events.

## Introduction

We have developed a novel biosensing device capable of detecting the real-time, dynamic *in vivo* oxidative status (IVOS) in living systems^1^. The sensor creates a microenvironment capable of detecting macromolecules containing carbonyl groups, which accumulate as a result of carbonylation in the presence of cytotoxic lipid peroxidation products such as aldehydes. As lipid peroxidation can occur non-enzymatically when reactive species interact with intra- and intercellular membranes, the incidence of macromolecule carbonylation serves as an indicator of cellular dysfunction and an oxidative steady-state typical of diseased or compromised cells^2^. Initial studies validated the IVOS biological sensor by demonstrating carbonylation detection using a beta-hydroxybutyrate dose response and measurements in anesthetised mice and pig tissues^1^. While the IVOS sensor allows for sensitive, site specific detection of oxidative stress, further disease and species-specific characterisation and validation, such as in cystic fibrosis (CF), are required to establish its utility.

Reactive species, including ROS and nitrogen derived reactive species, represent a critical signalling mechanism that regulates cellular activities, but their presence is generally transient to prevent accumulation. Cells have multiple mechanisms – including the ER and mitochondrial UPR as well as modification of antioxidant capacity – to maintain reactive species within a homeostatic range^2^. Excessive concentrations of reactive species result in an oxidative environment that decreases organelle function, induces calcium dysregulation, and increases the risk of cellular apoptosis. Collectively these conditions are referred to as oxidative stress^3,4^. Oxidative stress is a complex process with a multitude of metabolic processes involved, and even more signaling pathways affected.

CF is a chronic, progressive and debilitating inherited condition that affects multiple organs, especially those with epithelial tissue including airways, intestines, oesophagus, and reproductive organs^5,6^. CF arises due to pathogenic variants of the Cystic Fibrosis Transmembrane Conductance Regulator (*CFTR*) gene. *CFTR* dysfunction results in altered CFTR-dependent chloride and bicarbonate transport and dysregulation of the epithelial sodium channel (ENaC) activity. The resulting ion imbalance leads to dehydration of the epithelial surfaces, and in the lungs this results in accumulation of viscous mucus^7–9^. These conditions make CF lungs vulnerable to infection, resulting in further inflammation, injury and impaired function^10,11^.

The most common mutation is *Phe508del*, which is designated as a class II *CFTR* mutation. Class II mutations produce incorrectly folded proteins that cannot be shuttled to the cell membrane and remain in the endoplasmic reticulum (ER) awaiting degradation^12^. As this protein builds up in the ER, reactive oxygen species (ROS) production increases and stress pathways such as the unfolded protein response (UPR) are activated^13^. The combination of protein accumulation, infection and subsequent inflammatory responses chronically stimulate ER stress and the UPR. This generates a perpetually oxidative environment with decreased antioxidant capacity within CF epithelial tissues compared to wildtype^14^. As ROS interacts with cellular components, organelle function decreases, inducing further oxidative stress conditions within the cell^15^.

Elevated oxidative stress (OS) levels have been observed in both CF animal models and humans, where these conditions are associated with excessive neutrophil-driven inflammatory responses that contribute to fibrotic tissue formation^13,16^. As such, numerous clinical and preclinical studies suggest that elevated oxidative stress, coupled with the pronounced pro-inflammatory milieu observed in CF phenotypes, likely exacerbates susceptibility to recurrent infection–inflammation cycles^17^. Clinical interventions have targeted ER stress but with varied success.

While large animal models such as pigs and ferrets recapitulate the CF lung phenotype observed in humans, their use raises animal welfare concerns due to the disease severity, which also entails significant resource demands. Rodent models offer distinct advantages for investigating the mechanisms associated with CF as they do not display the severe disease phenotypes that compromise animal welfare. This makes them easier to breed and house compared to larger animals, thus allowing for larger study sample sizes^18^. We previously developed *Phe508del* and *CFTR* knockout CF rat models which, as with other rodent models, lack overt lung disease, with no histological changes consistent with a mucus plugging phenotype observed in the lung tissues^19^. However, using highly sensitive real-time X-ray Velocimetry ventilation imaging techniques combined with flexiVent lung mechanics assessments, we have observed statistically significant differences in lung function compared to wildtype^20^. In addition, we identified differences in the molecular profile of CF rats lungs indicating that airway epithelial cells exist in partial epithelial-mesenchymal transition (EMT) states and exhibit increased Type 1 collagen and vimentin expression^21^. These findings suggest CF rats have elevated lung tissue remodeling despite the absence of obvious histological changes. As such, they serve as suitable models to study aspects of the CF phenotype observed in humans.

The aim of this pilot study was to utilise the IVOS sensor to characterise oxidative environmental differences in the CF *Phe508del* and CFTR knockout rat models. To achieve this the IVOS biosensor was applied to epithelial tissues from wildtype, *Phe508del*, and *CFTR* knockout CF rats to determine whether changes in oxidative status can be detected.

## Methods

### Rats

This project was conducted under the approval of The University of Adelaide Animal Ethics Committee (M-2024-021). Wildtype (n = 10; 6 female, 4 male), *Phe508del* (n = 8; 4 female, 4 male) and knockout (n = 6; 5 female, 1 male) Sprague-Dawley rats from the Cystic Fibrosis Airway Group CF breeding colonies were obtained via scavenging excess animals immediately post-mortem^21^. Rats ranged in age from 54-374 days with a mean of 143.4 days and median of 109 days.

### IVOS sensor

The IVOS sensor was manufactured using previously published methodology^12^. Briefly, the tip of a glass core fibre (Thorlabs; M29L05) was dipped into a sensor-polymer composite material for 1 s. The fibre tip was polymerised to form a solid bead at the end of the fibre and dip cleaned using a solution of a 50/50 ratio of acetone and isopropanol, followed by a solution of isopropanol only. The functionalised fibre was coupled to a laser light source (Integrated Optics MatchBox CW) that emitted pulses of excitation light (488 nm) to the sensor tip via a bifurcated fibre bundle (Thorlabs; RP21). Emission wavelengths were coupled back into the bifurcated fibre bundle, passed through a 488 nm long-pass edge filter (Semrock; BLP01-488R-25) and characterised using an Ocean Optics QE Pro spectrometer. Emission wavelengths between 515 and 520 nm were used for statistical analysis.

### Measurements of epithelial tissues

Rats were humanely killed using CO_2_ asphyxiation. Death was confirmed through absence of heartbeat and ocular relaxation. Within two minutes of confirmed death, the IVOS sensor was used to collect measurements from various luminal epithelial tissues (Figure 1). Rats were hung by their teeth and the sensor placed down the oesophagus (Figure 1A). A tracheotomy was then performed, allowing access to insert the sensor into the trachea (Figure 1B). The sensor was then moved into the lungs until pressure could be felt through stretching bronchi (Figure 1C) and then the sensor was retracted for a relaxed bronchus (Figure 1D). The head of the rat was then propped up to allow the insertion into a nasal airway for the upper mucosal airways (Figure 1E). Finally, the sensor was inserted into the reproductive tract (cervical vaginal mucosa in females and preputial mucosa in males (Figure 1F). A minimum of 20 technical IVOS replicate measurements were performed for each tissue type and per animal. Each replicate consisted of excitation (488 nm) from the optical fibre for four sec, followed by retrieval of the emission spectra (515-520 nm)^1^. The sensor was submerged in 0.9% saline between animals. Different metrics were used to measure the oxidative environment (Table 1).

**Figure 1:**
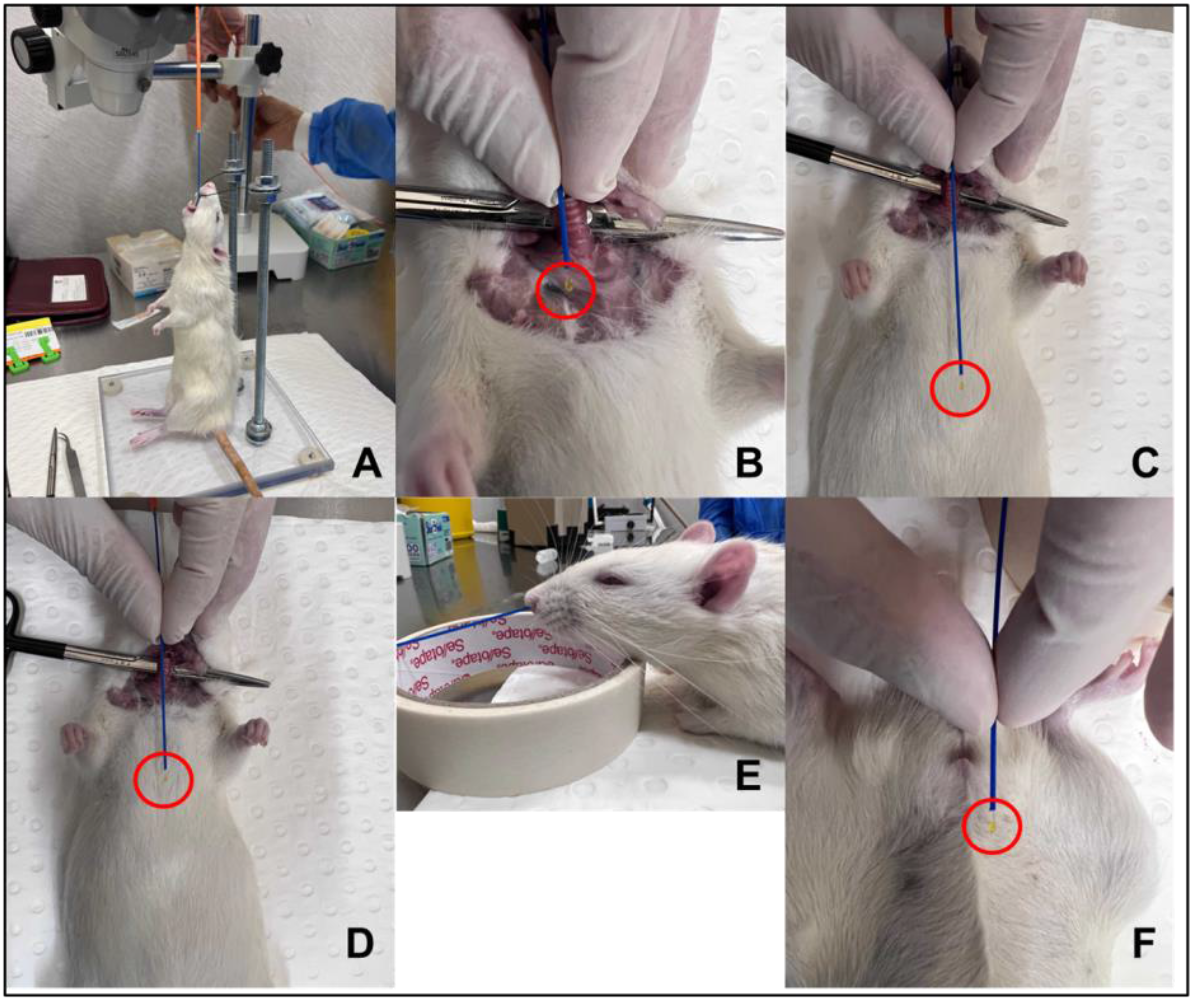
Representations of the location the IVOS sensor **(red circle)** was placed for measurements in the **A)** oesophagus, **B)** trachea, **C)** stretched bronchus, **D)** relaxed bronchus, **E)** upper nasal mucosa, and **F)** reproductive organs. The sensor is shown externally of the rat to indicate how far in the sensor was placed.

**Table 1:** IVOS metric definitions.

| Measurement | Definition |
| --- | --- |
| Arbitrary IVOS units (ArbU) | Raw baseline fluorescence measurements produced by the IVOS |
| Normalised arbitrary IVOS units (Normalised ArbU) | Arbitrary IVOS units were normalised to fluorescence levels when the probe was submerged in 0.9% saline. |
| Root Mean Squared of Successive Differences (RMSSD) | Quantifies how much the normalised ArbU changes from one replicate to the next, on average. |
| Ratio of RMSSD : normalised arbitrary IVOS units (RMSSD : normalised ArbU) | The RMSSD is calculated by accounting for the baseline oxidation (level Normalised arbitrary IVOS units) |

### Statistics

All data visualisation and statistical analysis was performed in GraphPad Prism (Version 10.1.0) software. Data are presented as mean ± SEM. One-way ANOVA and Tukey’s post hoc tests were used to compare IVOS measurements between different tissue types, as well as for the same tissues between wildtype, *Phe508del* and *CFTR* knockout rats. Significance was set at *p* < 0.05.

## Results

Measurements were collected using the IVOS from cervical-oesophagus, mid-trachea, stretched and relaxed bronchus, upper airway mucosa (ventral nasal passage), and reproductive organs (prepuce and cervical vaginal mucosa) of wildtype, Phe508del, and *CFTR* knockout rats. The data was then grouped and analysed by genotype to assess differences between different tissues (Figure 2), and by tissue type to determine differences between the genotypes (Figure 3).

**Figure 2:**
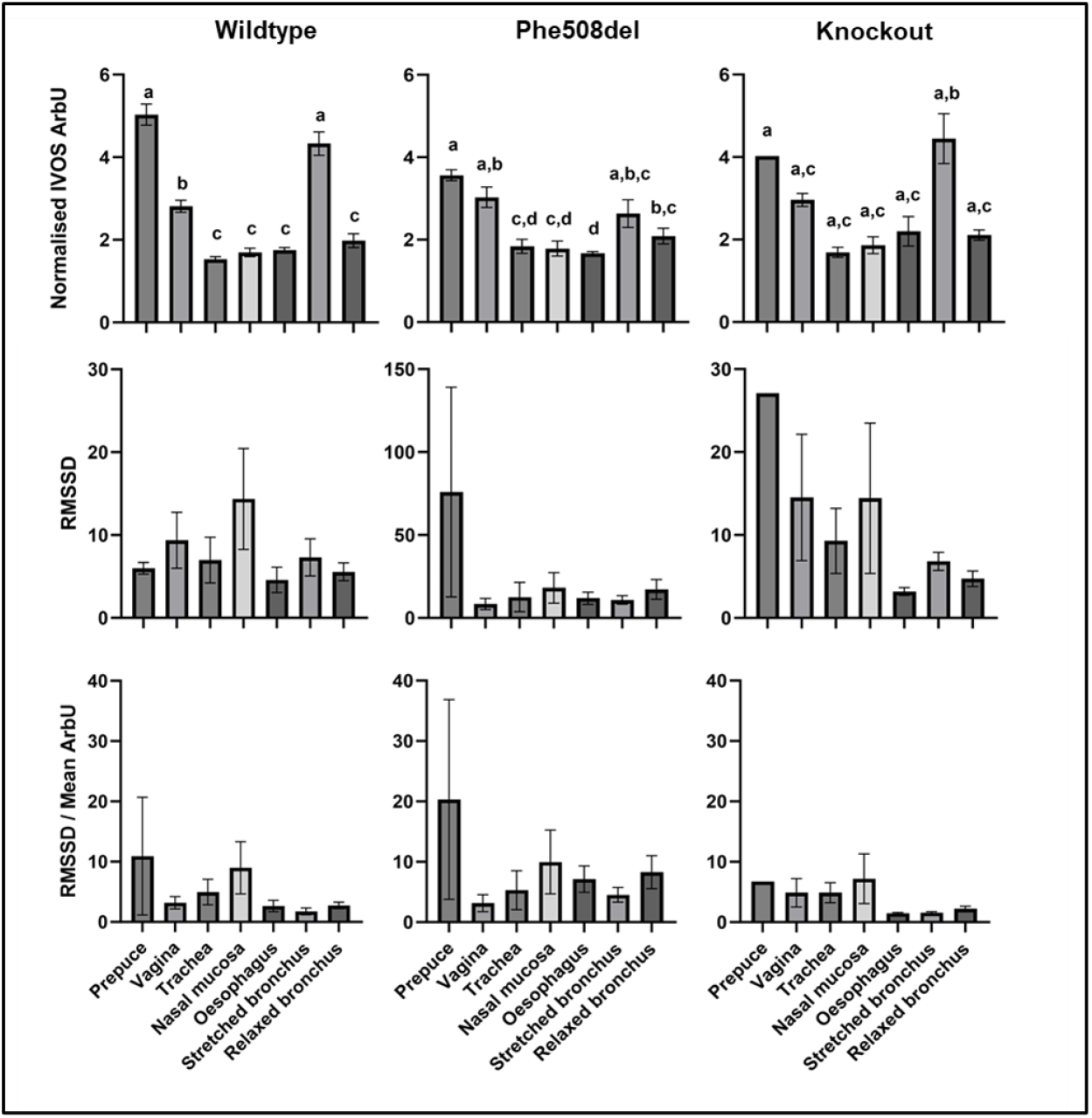
IVOS measurements in wildtype, *Phe508del* and knockout rat epithelium. Measurements were collected from the prepuce, vagina, trachea, nasal mucosa, oesophagus, stretched bronchus, and relaxed bronchus. The mean normalised IVOS ArbU, RMSSD, and RMSSD/ ArbU were determined. Results are presented as the mean ± SEM. One way ANOVA and Tukey’s post hoc test were carried out to determine significant differences. Groups that do not share a letter are significantly different from one another (p ≤ 0.05). Groups sharing at least one letter are not significantly different.

**Figure 3:**
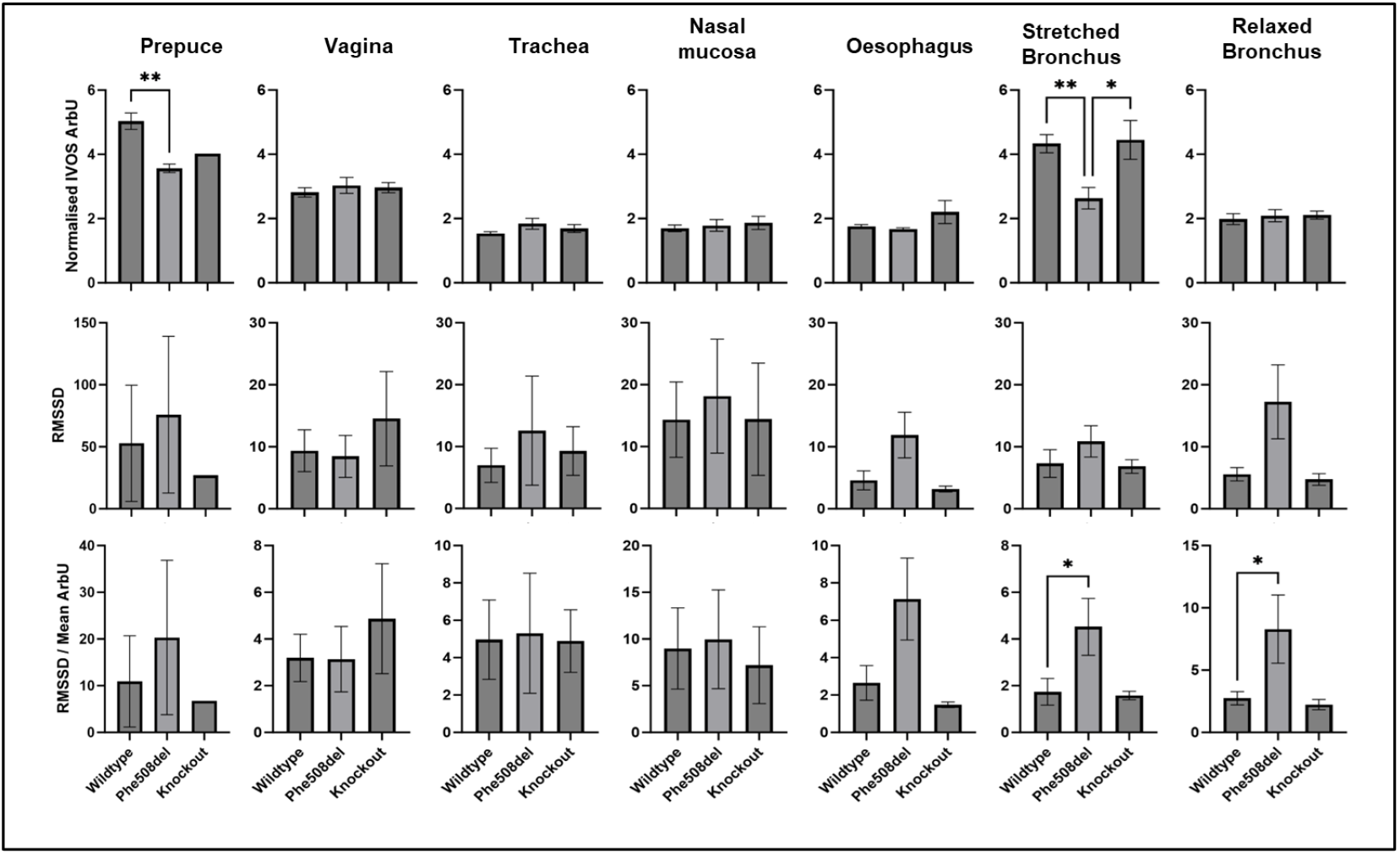
IVOS measurement comparisons of wildtype, *Phe508del* and knockout rats for each tested epithelium. The mean normalised IVOS ArbU, RMSSD, and RMSSD/ Normalised ArbU were determined for the prepuce, vagina, trachea, nasal mucosa, oesophagus, stretched bronchus, and relaxed bronchus. Results are presented as the mean ± SEM. One way ANOVA and Tukey’s post hoc test were carried out to determine significant differences, represented by *, with significance set to *p* ≤ 0.05.

## Baseline level of IVOS ArbU within each rat genotype varies significantly depending on tissue type

We determined the baseline arbitrary IVOS units of each epithelial tissue of wildtype, *Phe508del* and knockout CF rats. In wildtype and *Phe508del* rats, the highest IVOS ArbU measurements were observed in the prepuce and stretched bronchi, while trachea, nasal mucosa, oesophagus, and relaxed bronchi all displayed the lowest levels (Figure 2). The results from knockout rats were similar, however due to tissue availability, preputial measurements were performed on a single male rat. The stretched bronchus demonstrated markedly elevated normalised IVOS ArbU values compared to all other tissues assessed.

Despite differences in IVOS ArbU across the three genotypes, we did not see any significant differences in the RMSSD or the ratio of RMSSD to ArbU.

### Oesophagus and bronchus in *Phe508del* rats have greater IVOS fluctuations than wildtype and CFTR-Knockout rats

We then assessed the differences between wildtype, Phe508del, and knockout rats in each tissue (Figure 3). The normalised IVOS ArbU of the prepuce of *Phe508del* rats was lower compared to wildtype rats (p = 0.0052). Similarly, the normalised IVOS ArbU of the stretched bronchus of *Phe508del* rats was reduced compared to wildtype rats (p = 0.0088) and knockout rats (p = 0.0140). No differences were detected in the epithelial tissues of vagina, trachea, nasal mucosa, oesophagus, and relaxed bronchus.

There was an increase in the RMSSD for *Phe508del* rats for the oesophagus, stretched bronchus, and relaxed bronchus, however none reached significance compared to wildtype and knockout rats. To take the baseline differences in the IVOS ArbU into consideration, we then looked at the ratio of RMSSD to ArbU. Similarly to the RMSSD, an increase was observed in *Phe508del* esophagus, stretched bronchus, and relaxed bronchus, with a significant difference in both the stretched and relaxed bronchus compared to wildtype (p = 0.0493; p = 0.0489, respectively). No differences in the RMSSD and the ratio of RMSSD and ArbU were observed for prepuce, vagina, trachea, and nasal mucosa.

## Discussion

Oxidative stress is linked to worse health outcomes in people with CF (pwCF)^16^, while timely modulation of oxidative status would likely improve health outcomes. Thus, the capability to measure oxidative status in real time, and in specific regions and tissues, has the potential to drastically improve diagnostic times, the course of treatment, and the success of interventions. In this study we assessed *in vivo* differences in oxidative status using the IVOS biosensor in two CF rat models for the first time. We determined that various epithelial tissues show different basal levels of oxidative status and observed that *Phe508del* rats have greater IVOS fluctuations than wildtype and knockout rats.

The stretched bronchus recorded significantly higher IVOS ArbU compared to the relaxed bronchus in wildtype and knockout rats, suggesting increased oxidative stress induced by or during the stretching motion. Conversely, the stretched bronchus in *Phe508del* rats displayed reduced mean fluorescence but an increased RMSSD/mean ratio compared with wildtype rats, indicating a lower steady-state oxidative environment but large transient oxidative events. This pattern could be due to baseline antioxidant buffering coupled with episodic ROS bursts, potentially arising from *Phe508del*-associated ER stress. Studies have shown that stretched cells or tissue results in increased ROS production. Stretching of the retinal cell line ARPE-19 for 15 minutes was sufficient to significantly increase ROS formation^22^. Mechanical stretching can also induce mitochondrial driven apoptosis and accumulation of pro-apoptotic factors in cardiomyocytes^23^. Notably, the other two tissues tested that required some level of navigation and tissue extension to insert the probe were the prepuce and cervical region of the vagina, which also demonstrated increases in IVOS measurements when compared to tissues assessed with minimal manipulation (Fig. 2). Further investigation of this phenomenon is required, but tissue stretch could be an important variable to control for future studies. Ultimately, whether there are basal differences between the organs is also worth further investigation in larger, well powered studies.

Other mechanisms that contribute to CF disease progression may also increase oxidative status markers that are measurable with the IVOS sensor, including immune responses and epithelial mesenchymal transition (EMT)^21^. Both of these mechanisms are associated with ROS production, as stimulated neutrophils produce a variety of ROS, and ROS directly activate EMT pathways^24,25^. Therefore, it is possible that excessive production of ROS through neutrophil activity or ROS accumulation that upregulates EMT could also subsequently increase carbonylation of local macromolecules in a manner that is measurable with the IVOS sensor. It would be of interest to further investigate both the immune response and EMT mechanisms from CF models in conjunction with IVOS technology.

Although this is the first study to assess IVOS measurements in an animal model of disease, there are some limitations to this study. Freshly killed rats likely have physiological differences to live rats that could alter measurements. After death, the redox balance shifts to a more oxidative environment due to less antioxidant production but increased cellular damage^26^. Future studies should be conducted on live animals. A second limitation is that CF rats have relatively healthy respiratory tracts, with only bioelectric defects in the nasal epithelium observed^19,20,27^. Unlike in pwCF, there is no increased inflammation or bacterial burden compared to wildtype rats. Inflammation and bacterial burden is likely to be a strong contributing factor to an oxidising environment. Some studies have shown that CF lungs fail to modulate lipopolysaccharide (LPS) or *Pseudomonas aeruginosa* induced inflammation as quickly as wild type in a mouse model^11^. Therefore, studies that induce an inflammatory response should also be investigated with the IVOS. A third limitation of this study was the inability to accurately categorise ROS present within the cells. The IVOS technology can be used as an indicator of oxidative stress by measuring the level of macromolecule carbonylation, which occurs through chain reactions originating from reactive species interactions with cellular components. As such, IVOS is unable to discriminate between the different types of reactive species but rather is used as an indication of cellular oxidation conditions or oxidative status by measuring processes secondary to ROS accumulation. Although it will provide overall oxidative stress levels, more detail such as which species are actively produced will still require molecular profiling, especially for studies investigating immune responses and EMT.

## Conclusion

In conclusion, the IVOS technology was able to measure differences in carbonylation between wildtype and CF rats by using the fluorescent probe in different epithelial tissues. Although we were only able to find differences in the mean fluorescence between wildtype and Phe508del in the stretched bronchus, the RMSSD and the ratio of RMSSD to mean fluorescence does appear to be more informative in identifying at least short-term fluctuations in the redox balance. Furthermore, these changes appear to be specific to the class II Phe508del mutation, which is known to result in an oxidative environment due to the ER retention of the misfolded Phe508del CFTR protein, opposed to the absence of CFTR protein in the knockout model. This is the first study to use IVOS assessment of a genetic disease with known oxidative stress link and warrants further analysis.

## Notes

### Competing Interest Statement

The research presented involves technology developed at the University of Adelaide, and is now associated with Sensrz Pty Lid. R. Kostecki is a Founder and Managing Director at Sensrz Pty Ltd.

